# Toxic Benthic Filamentous Cyanobacteria in Lakes and Rivers of South-Central Quebec, Canada

**DOI:** 10.1101/580035

**Authors:** Barry Husk, Debra Nieuwenhuis

## Abstract

Toxic cyanobacteria are a present and growing threat to ecosystems and public health worldwide. However, most research and regulatory measures have focussed on the planktonic forms of cyanobacteria, with consequently little information available concerning potentially toxic benthic filamentous forms. Through a regional study of ten lake and river sites, including some which are sources of municipal drinking water, this investigation confirms for the first time the widespread presence of potentially toxic benthic filamentous cyanobacteria in south-central Quebec. These findings indicate that water quality monitoring programs in this region need to consider benthic cyanobacteria as a potential source of toxins.

## Discussion

Toxic cyanobacteria are increasingly present in aquatic environments worldwide, causing ecosystem impairment, as well as threatening public health (Otten & Paerl 2015). In addition, continued aquatic eutrophication and global warming are expected to exacerbate this situation, causing harmful cyanobacterial algal blooms to intensify in frequency and duration (Mantzouki *et al.* 2018). Cyanobacteria occur in freshwater environments in several forms, notably within the water column (planktonic forms), or attached to substrates (benthic forms). However, studies of ecological and health risks, as well as regulatory measures associated with cyanotoxins, have been primarily related to planktonic species. Consequently, relatively little is known about benthic cyanobacteria populations (Quiblier *et al.* 2013). It is, however, well-established that benthic cyanobacteria are producers of cyanotoxins, including hepatotoxic microcystins, nodularins and cylindrospermopsins; neurotoxic saxitoxins, anatoxin-a and homoanatoxin-a; as well as dermatotoxins, such as lyngbyatoxin (Bernard *et al.* 2017). Depending on the class of cyanotoxin, these can create potential threats to human health by exposure through drinking water or through direct contact. In addition to human health risks, cyanotoxins produced by benthic species can present threats to both wild and domestic animals. In particular, numerous accounts of benthic cyanotoxin poisoning and deaths of dogs have been recorded, including in Canada (Hoff *et al.* 2007), and elsewhere (Cadel-Six *et al.* 2007). In addition to producing cyanotoxins, certain species of filamentous benthic cyanobacteria are also known to harbour fecal indicator bacteria, including *Escherichia coli,* potentially further impacting human health and water quality (Vijayavel *et al.* 2013). As a result, more study concerning the presence and ecology of potentially toxic benthic cyanobacteria is recommended by public health authorities (Government of Canada 2016).

Some of the most significant climate-related increases in cyanobacteria are projected to occur in the north-eastern region of North America (Chapra *et al.* 2017). Located in this area, the relatively populated sub-region of south-central Quebec has a concentration of lakes and rivers, many of which are sources of municipal drinking water. It is also a popular area for tourism and recreational use of waterways, all of which create opportunities for exposure to cyanotoxins. The persistent presence of the planktonic form of cyanobacteria in the many lakes of this region is also well known (Government of Quebec 2019). However, knowledge related to the presence of benthic filamentous cyanobacteria in this region is virtually non-existent and limited primarily to studies focused on the fluvial Lake St. Pierre (St. Lawrence River), where the benthic cyanobacteria species *Lyngbya wollei* has been studied (Hudon *et al.* 2014). There are currently no norms or government monitoring programs examining the presence or impacts of benthic cyanobacteria in this region.

To respond to this research gap, as well as the expressed desire of public health authorities to better understand the ecology and potential problems posed by these species, this study has examined, for the first time on a regional scale in this area, whether potentially toxic benthic cyanobacteria are present in certain surface waters of this region, including some which are sources of municipal drinking water.

A total of 10 sites were chosen in south-central Quebec, including both lakes and rivers, based on their use either as a site for recreational activities, and/or a source of municipal drinking water (Table 1). These sites are located within the surface watersheds of either the St. Francis or the Yamaska Rivers. Sampling was conducted between the end of May and the beginning of August, 2018, in the littoral zones of aquatic areas accessible for recreational purposes. At each site approximately 100 metres of shoreline was visually surveyed for the presence or accumulation of benthic cyanobacteria. Cyanobacteria were captured directly with, and stored in, high density polyethylene (HDPE) screw-cap bottles, along with water from the sampling site. Lugol’s solution was added at 0.2% and samples were transported on ice, stored at 4°C, and shipped overnight for taxonomic analysis (Water’s Edge Scientific, LLC, Baraboo, WI, USA). This analysis was performed by light microscopy (Motic Microscopes, model BA310) and identification was made to genus and species using standard taxonomic references (Komárek 2013; Komárek & Anagnostidis 2005). The results of these findings are described in Table 1. All species examined were potentially cyanotoxin-producing species. Figure 1 illustrates photomicrographs of certain samples of species discovered in this study.

**Table 1.** Summary of benthic cyanobacteria taxa, the site location and municipality of samples, whether these sites were a source of municipal drinking water, and potential cyanotoxins produced by the taxa present.

| Site | Municipality | DW<sup>1</sup> | Taxa | Potential cyanotoxins<sup>2</sup> |
| --- | --- | --- | --- | --- |
| Petit Lac St. François | St-F-X-de-Brompton |  | *Lyngbya wollei* | C, S |
| Petit Lac St. François | St-F-X-de-Brompton |  | *Scytonema crispum* | S |
| St. Francis River, Bellevue Park | Drummondville | * | *Oscillatoria princeps* | A-a |
| St. Francis River, St. Thérèse Park | Drummondville | * | *Oscillatoria princeps* | A-a |
| Lake Boivin, Lemieux Reservoir | Granby | * | *Phormidium sp.* | A-a, HA-a |
| Lake Boivin, Lemieux Reservoir | Granby | * | *Oscillatoria sp.* | A-a, HA-a, C |
| Lake Boivin, Lemieux Reservoir | Granby | * | *Planktothrix sp.* | M |
| St. Francis River | Pierreville | * | *Phormidium sp.* | A-a, HA-a, M |
| Watopeka River | Windsor | * | *Lyngbya sp.* | C, S |
| Watopeka River | Windsor | * | *Oscillatoria sp.* | A-a, HA-a, C |
| Watopeka River | Windsor | * | *Microcoleus sp.* | A-a |
| Lake Memphremagog, Fitch Bay | Stanstead | * | *Oscillatoria limosa* | A-a, HA-a, M |
| Lake Bromont | Bromont |  | *Lyngbya sp.* | C, S |
| Lake Bromont | Bromont |  | *Oscillatoria sp.* | A-a, HA-a, C |
| Lake Davignon | Cowansville | * | *Lyngbya wollei* | C, S |
| Magog River, Lake des Nations | Sherbrooke | * | *Lyngbya wollei* | C, S |
| Magog River, Lake des Nations | Sherbrooke | * | *Oscillatoria sp.* | A-a, HA-a, C |

<sup>1</sup>Source of municipal drinking water
<sup>2</sup>A-a/Anatoxin-a, HA-a/Homo-anatoxin-a, C/Cylindrospermopsin, M/Microcystin, S/Saxitoxin (Bernard et al., 2017)

**Figure 1.**
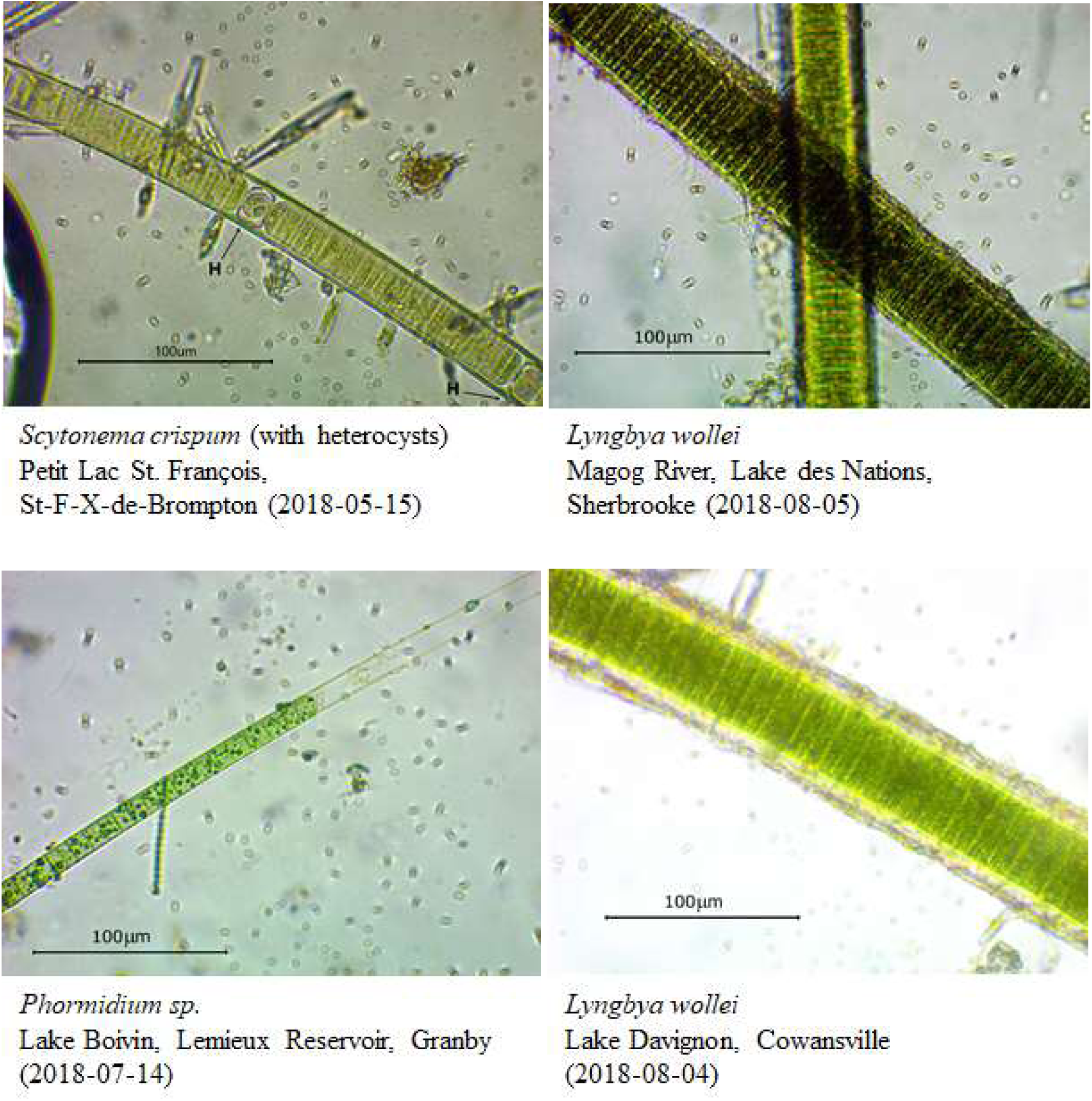
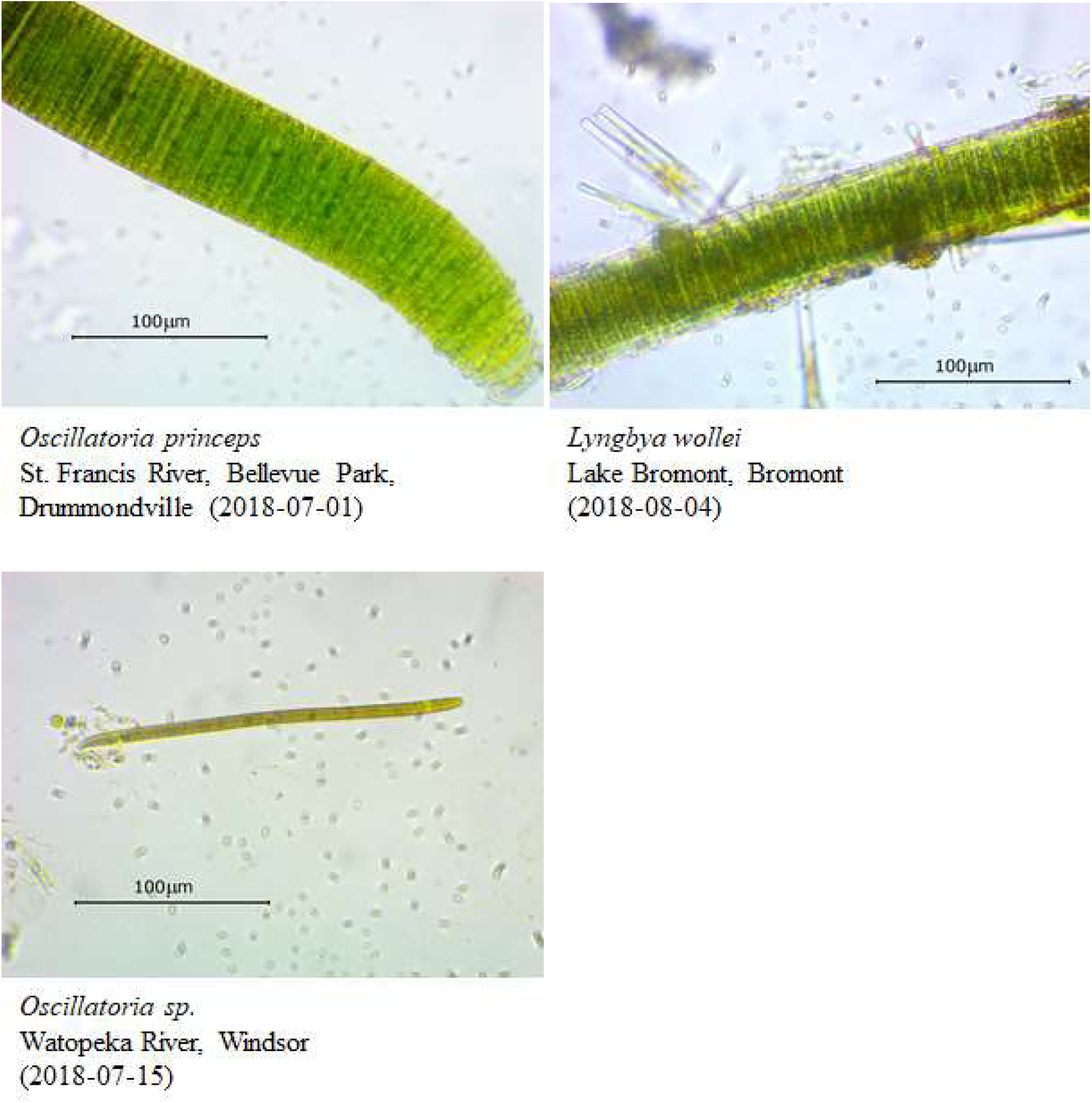
Photomicrographs of representative benthic cyanobacteria genera examined in this study. Scale bars = 100 μm.

## Conclusions

Improved understanding of benthic cyanobacterial ecology is needed in order to identify and predict where and when such cyanobacterial proliferations may negatively affect aquatic ecosystems and human health. This study contributes to filling an important research gap by confirming the presence of potentially toxic benthic filamentous cyanobacteria in lakes and rivers of south-central Quebec, including some which are sources of drinking water. The findings further indicate that water quality monitoring programs need to consider benthic cyanobacteria as a potential source of toxins, as well as highlighting an additional freshwater habitat where cyanobacteria need to be monitored. Using these findings as a basis for further studies, we call on researchers, public health officials and regulators to further examine this subject, including the toxicity of these species.

